# Facing the heat: thermoregulation and behaviour of lowland species of a cold-dwelling butterfly genus, *Erebia*

**DOI:** 10.1101/024265

**Authors:** Irena Kleckova, Jan Klecka

## Abstract

Understanding the potential of animals to immediately respond to changing temperatures is imperative for predicting the effects of climate change on biodiversity. Ectothermic animals, such as insects, use behavioural thermoregulation to keep their body temperature within suitable limits. It may be particularly important at warm margins of species occurrence, where populations are sensitive to increasing air temperatures. In the field, we studied thermal requirements and behavioural thermoregulation in low-altitude populations of the Satyrinae butterflies *Erebia aethiops*, *E. euryale* and *E. medusa*. We compared the relationship of individual body temperature with air and microhabitat temperatures for the low-altitude *Erebia* species to our data on seven mountain species, including a high-altitude population of *E. euryale*, studied in the Alps. We found that the grassland butterfly *E. medusa* was well adapted to the warm lowland climate and it was active under the highest air temperatures and kept the highest body temperature of all species. Contrarily, the woodland species, *E. aethiops* and a low-altitude population of *E. euryale*, kept lower body temperatures and did not search for warm microclimates as much as other species. Furthermore, temperature-dependence of daily activities also differed between the three low-altitude and the mountain species. Lastly, the different responses to ambient temperature between the low- and high-altitude populations of *E. euryale* suggest possible local adaptations to different climates. We highlight the importance of habitat heterogeneity for long-term species survival, because it is expected to buffer climate change consequences by providing a variety of microclimates, which can be actively explored by adults. Alpine species can take advantage of warm microclimates, while low-altitude grassland species may retreat to colder microhabitats to escape heat, if needed. However, we conclude that lowland populations of woodland species may be more severely threatened by climate warming because of the unavailability of relatively colder microclimates.

## Introduction

Ongoing climate changes induce range shifts of many animals and plants [1,2], however species responses to climate change are individualistic. At the population level, they adapt by evolutionary changes [3–5]. These adaptive responses to environmental changes contribute to the demarcation of current species distribution ranges and also induce diversification. Selective pressure of large environmental changes leads to shifts of allele frequencies of genes responsible for physiological functions [3], and creates a phylogenetic signal in genus-level phylogenies [6, 7]. At the individual level, animals acclimate [8], adjust their phenology [9], and use a wide repertoire of behaviours to regulate their body temperature [4, 10].

Behavioural plasticity of ectotherms enables them to keep their body temperature at the optimal level and thus to optimize functioning of their physiological processes at short time scales [10]. Behavioural thermoregulation works mainly by changes in the timing of daily activities or exploration of various (micro) habitats [11–13]. The ability to use locally suitable microhabitats facilitates species survival under ongoing climate change [10]. Ectotherms search for warmer microclimates at the cold margins [12, 14] and decrease activity or search for shade at the warm margins of occurrence [15–18]. The comparison of species responses at contrasting climatic range margins, such as low-vs. high-altitude margins, may empower effective targeting of conservation activities, especially in declining species with limited potential for range shifts [2, 19, 20].

Despite the great behavioural plasticity of many species, responses to climate change may be limited by various constraints. For example, tropical lizards of forest interiors suffer from a higher risk of overheating than their open-habitat congeners [21]. Under a climate warming scenario, open-habitat species have the potential to explore novel habitat types in the forest interior by adjusting their microhabitat use while forest-dwellers have little chance of doing so because they already occupy the coolest available microhabitats [21]. Despite many obvious advantages of behavioural plasticity as a rapid way to respond to environmental changes [10], behavioural thermoregulation could also limit physiological adaptation, which is necessary for species' long-term survival [8, 22, 23]. Whether this so-called ˮBogertˮ effect has a general validity is not yet clear, but it represents an important possibility, with implications for predicting climate change effects.

Organisms living at high altitudes were among the first to attract considerable attention to the issues of species survival under climate change because of apparent range shifts along altitudinal gradients [1, 24, 25]. However, recent studies report that lowland fauna seem to be even more sensitive to ongoing climate change than mountain species. Detailed studies of internal traits such as immune system [26], thermoregulation [21] or thermal limits [27] surprisingly report higher sensitivity of lowland species to increasing temperatures. Moreover, also extrinsic factors such as existence of extremely diverse (micro)habitats and topographies [2, 14, 28], or feasibility of uphill shifts along the altitudinal gradient [2], support long-term survival of species in the mountains. On the contrary, lowlands provide a more homogeneous environment; in addition, they are fully exposed to human activities such as agriculture, forestry and urbanization. In concordance, a large comparative study by Roth et al. [2] showed that mountain assemblages are more stable, i.e. less vulnerable, than lowland ones in relation to climate change.

Here, we focus on thermal ecology and microclimate utilisation at high- and low-altitude margins of occurrence in adult butterflies of the genus *Erebia.* Alpine *Erebia* species are well adapted to extreme high-altitude environments, where they are able to maximize activity during short periods of favourable weather and they do not seem to be at risk of overheating in a warming environment [14]. However, the situation in aberrant lowland species and populations of this mostly mountain genus is not known. Thus, we examined thermoregulatory strategies in adults of two lowland species at the upper thermal margins of their occurrence and of a low-altitude mountain population of a montane-belt species. We compared our findings with data on seven alpine species reported in [14]. We discuss the role of behavioural thermoregulation and physiological adaptations of lowland and mountain species of *Erebia* for dealing with climate change.

## Materials and Methods

### Study group and study sites

The genus *Erebia* Dalman, 1816 is a popular group in studies of eco-physiological adaptations [14, 29] and biogeography [30–32], as well a subject of increasing interest from the conservation perspective [14, 18, 33–36]. In our study, we compared thermoregulation of seven alpine species occurring at high altitudes in the European Alps (see [14] for more details) with thermoregulation of a low-altitude population of a mountain species *E. euryale* (Esper, 1805), and two aberrant lowland species, *E. aethiops* (Esper, 1777) and *E. medusa* (Fabricius, 1787). No specific permissions were required for this project because it did not involve any protected species and the field work took place outside of any protected areas. The field work was carried out on public land, hence, no access permissions were needed.

#### Mountain species

The seven alpine species *Erebia alberganus* (Prunner, 1798), *E. euryale* (Esper, 1805), *E. ligea* (Linnaeus, 1758), *E.melampus* (Fuessly, 1775), *E. montana* (de Prunner, 1798), *E. pandrose* (Borkhausen, 1788) and *E. cassioides* (Reiner and Hohenwarth, 1792) were studied in the Austrian Alps, Tirol, close to the town of Sölden, at an altitude of 1500-1800 m a.s.l. (see [14] for more details). These species occur in montane and alpine vegetation belts [37]. Another, low-altitude, population of *E. euryale* was studied in the Šumava Mts., Czech Republic, close to Borová Lada village, in the area of a former village, Zahrádky (13.68032°E,48.97362°Ν, altitude 930 m a.s.l.). *E. euryale* inhabits clearings and road margins within spruce forest in the area and was frequently observed nectaring on nearby pastures. The species *E. euryale* was thus represented by two populations: by a high-altitude population from the Austrian Alps (*E. euryale*-Alps) and by a low-altitude population from the Šumava Mts., Czech Republic (*E. euryale*-CZ).

#### Lowland species

*E. aethiops* and *E. medusa* inhabit a rather wide range of altitudes from lowlands to sub-alpine zones [37]. Both lowland species originated during Miocene diversification of a European clade of the genus *Erebia* and they are members of a phylogenetically distinct species group [32]. Their conservation status is Least Concern for Europe [38], but both species have experienced local declines throughout their ranges [39, 40] attributed to habitat alteration [18, 36]. Moreover, both species are hypothesized to be negatively affected by ongoing climate change [17, 41]. We studied their lowland populations in an ex-military area, Vyšný, in the vicinity of the town of Český Krumlov, SW Czech Republic (48°49’N, 14°1’E, altitude 550 m a.s.l.). The area includes a mosaic of dry grasslands, shrubs and woodlands. Both species co-occur in this relatively warm and dry area, probably on their upper thermal limits. *E. aethiops* (flight period: mid-June to August) is a woodland species, which inhabits forest edges, small grassland patches and sparse woodlands within the area [17, 18], while *E. medusa* (flight period: May to mid-July) is found in more open habitats - xeric grasslands with scattered shrubs.

### Measurements of body, microhabitat and air temperatures

Butterfly body temperature T_b_ was measured by a hypodermic micro-needle probe (0.3 mm in diameter) within 5 seconds after capture and values were recorded on Physitemp thermometer (model BAT-12) during August 2010 (*E. aethiops*), May 2012 (*E. medusa*), July 2012 (*E. aethiops, E. euryale-*CZ) and August 2012 (*E. euryale-*CZ). The alpine species were measured during their flight periods in 2010 and 2011 (see [14]). Butterflies were collected by a net during random walks within the study area during the entire daytime activity period (between 9 a.m. and 6 p.m.). During measurement, the microprobe was inserted into a butterfly’s thorax; the butterfly and the microprobe were shaded during the measurement. The same thermometer was used to measure microclimate *T_m_* and air *T_a_* temperatures. Microclimate temperature *T_m_* was measured immediately after *T_b_* measurement at the place where the butterfly was located before the capture (approximately 3 mm above substrate in the case of sitting or nectaring butterflies). Air temperature *T_a_* was measured at 1.5 m above ground by a shaded thermometer. Further, we classified individual behaviour prior to capture into three categories: sitting, nectaring and flight. All data on Czech butterflies are available in S1 File, data from Austria are available in [14] (Appendix A).

### Data analyses

The aim of our analyses was to compare thermoregulation of related butterflies occurring in mountains and in lowlands. All analyses were conducted in R 3.0.2 [42]. First, we tested the dependence of body temperature *T_b_* on microhabitat *T_m_* and air *T_a_* temperatures using Generalized Additive Models (GAMs) with cubic splines and maximal complexity of the fitted relationship set to *k* = 5 d.f. in mgcv 1.8-4 package for R [43]. The two relationships (*T_b_ ~T_a_* and *T_b_ ~T_m_*) were fitted for individual species (lowland and alpine populations of *E. euryale* were analysed separately). Data on alpine species reported by [14] were re-analysed the same way to facilitate comparison with the lowland species (the original conclusions of [14] were not affected).

We also searched for differences in active behavioural thermoregulation between lowland and mountain species. We estimated the difference between microhabitat and air temperatures *T_m_* — *T_a_* for settling and nectaring individuals (i.e. potentially actively thermoregulating) of each low-altitude and mountain species (separately for the two populations of *E. euryale*). Then, we used t-tests to test whether lowland species and *E. euryale* population also search for similarly warm microclimates as the alpine species [14].

Last, we tested how the proportion of settling, nectaring and flying individuals for individual species depends on air temperatures *T_a_* using GAMs with quasibinomial distribution. This analysis was conducted for the three types of behaviour and all lowland and mountain species/populations separately.

## Results

The two lowland species, *E. medusa* and *E. aethiops,* experienced higher maximal and higher minimal air temperatures *T_a_* than the seven mountain species studied (Fig. 1, Fig. 2A, Table 1). The low-altitude population of *E. euryale*-CZ experienced similar maximal air temperatures *T_a_* but ca 5°C higher minimal air temperatures *T_a_* compared to the alpine population of *E. euryale* (Fig. 2A). In spite of these differences in experienced air temperatures *T_a_,* experienced maximal microhabitat temperatures *T_m_* did not differ in species across different altitudes (Fig. 2B). Thus, alpine species were able to detect awarmer microclimate to compensate for the experienced low air temperatures (Fig. 3). Minimal microhabitat temperatures *T_m_* were higher in the lowland species corresponding to the generally warmer lowland environment.

**Figure 1.**
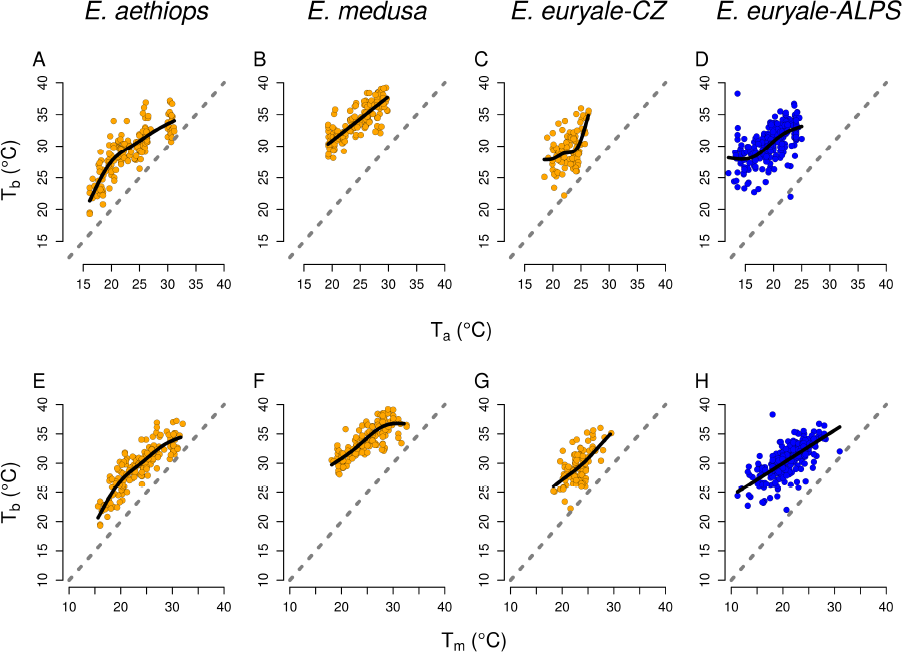
Body temperature depends on air and microhabitat temperatures. The relationship between body temperature *T_b_* and air temperature *T_a_* (A-D) and body temperature and microhabitat temperature *T_m_* (E-H) in two lowland species of *Erebia* butterflies and in low- (*E. euryale*-CZ) and high-altitude (*E. euryale*-Alps) populations of a mountain species, *E. euryale.* See Table 2 for detailed results of statistical tests.

**Table 1.** Overview of temperature data. Summary of measured values for the two lowland butterfly species *Erebia aethiops* and *E. medusa* and for low- and high-altitude populations of *E. euryale* are shown; data for the remaining alpine congeners are available in [14]. *N* (*F, M*) = the number of individuals measured (females, males); *T_a_* = air temperature; *T_m_* = microhabitat temperature; *T_b_* = body temperature.

| Species | Location | Flight period | $N(F, M)$ | | $T_a$ | $T_m$ | $T_b$ |
| --- | --- | --- | --- | --- | --- | --- | --- |
| *E. aethiops* | Czech Republic; 550 m a.s.l. | July - mid August | 161 (33, 128) | mean | 23.44 | 23.60 | 29.42 |
|  |  |  |  | median | 23.10 | 23.80 | 29.80 |
|  |  |  |  | range | 16.15-31.15 | 15.6-32.00 | 19.30-37.20 |
| *E. euryale*-CZ | Czech Republic; 930 m a.s.l. | July - mid August | 104 (33, 71) | mean | 22.80 | 23.09 | 29.36 |
|  |  |  |  | median | 23.05 | 23.30 | 29.05 |
|  |  |  |  | range | 18.80-26.35 | 18.30-29.50 | 22.20-36.00 |
| *E. euryale*-Alps | Austria; 1500-1800 m a.s.l. | July - mid August | 254 (79, 175) | mean | 19.67 | 21.02 | 30.57 |
|  |  |  |  | median | 20.00 | 21.00 | 30.95 |
|  |  |  |  | range | 12.00-25.00 | 11.20-31.00 | 22.00-38.30 |
| *E. medusa* | Czech Republic; 550 m a.s.l. | May - June | 154 (19, 135) | mean | 24.70 | 25.80 | 34.14 |
|  |  |  |  | median | 25.11 | 25.40 | 33.90 |
|  |  |  |  | range | 19.28-29.82 | 18.10-32.60 | 28.10-39.20 |

Body temperatures *T_b_* increased with slight deviations from linearity with increasing air temperatures *T_a_* as well as with increasing microhabitat temperatures *T_m_* in all species (Fig. 1, Fig. 2, Table 2). However, *E. euryale-CZ* and *E. aethiops* kept lower *T_b_* at similar *T_a_* than all other species, i.e. species from the Alps and also *E. medusa* (Fig. 2). Especially *E. aethiops* kept its body temperature only several C above the air temperature *T_a_* and had *T_b_* almost equal to *T_a_* at high *T_a_* values (Fig. 1). The lowland population of *E. euryale*-CZ and *E. aethiops* had a lesser tendency to heat up at given air temperatures *Ta* than the other species. *E. aethiops* slowly increased its *Tb* compared to *Ta* in a range of air temperatures 15-20°C and, after reaching the optimum around *Ta* = 21°C, the difference between its *Tb* and *Ta* decreased again (Fig. 2A; compare the fitted curve to the dashed line denoting *Tb* = *Ta).* It seems that *E. aethiops* thus has a narrow temperature range for activity compared to the other species. Contrary, *E. medusa* had thermoregulatory characteristics similar to the alpine species (Fig. 2). These species increased their *Tb* more at low than at high *Ta,* which suggests active thermoregulation by microclimate choice or body posture adjustment under low air temperatures.

**Figure 2.**
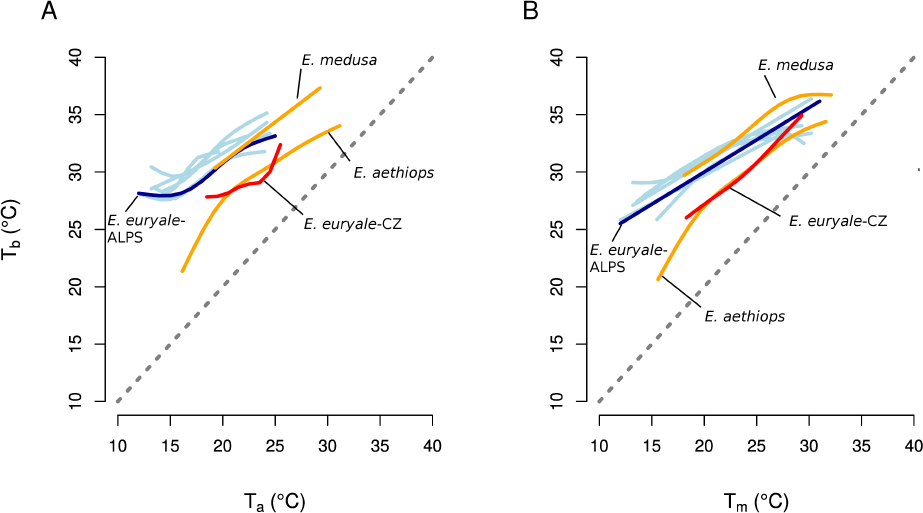
Comparison of body temperatures of lowland and alpine species and populations. The dependence of body temperature *T_b_* on air temperature *T_a_* (A) and on microhabitat temperature *T_m_* (B) of low-altitude (orange lines) and high-altitude (blue lines) species of *Erebia* butterflies fitted by generalized additive models. Two populations of *E. euryale* are shown in red (low-altitude) and dark blue (high-altitude). Only the fitted lines are shown to facilitate comparison. See Table 2 for detailed results of statistical tests.

**Table 2.**
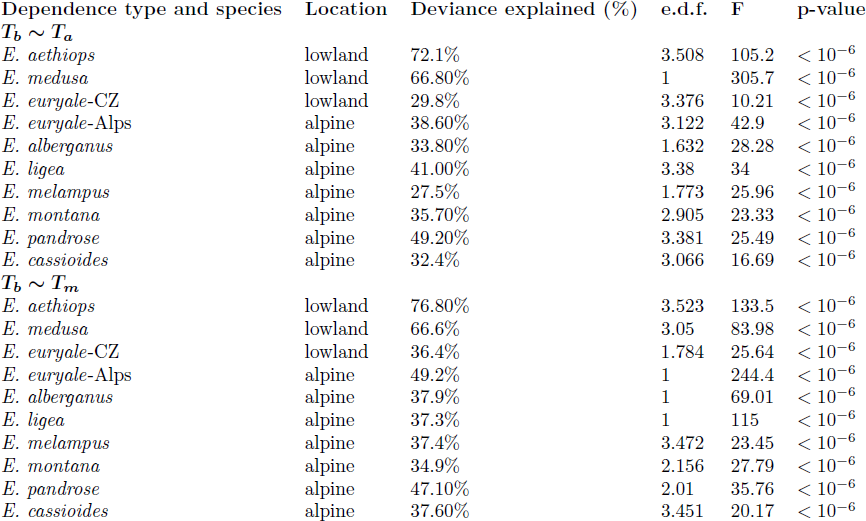
Body temperature depends on air and microhabitat temperatures. Results of generalized additive models (GAM) testing the dependence of body-to-air temperature excess *T_b_* — *T_a_* on air temperature *T_a_* and body-to-microhabitat temperature excess *T_b_* — *T_m_* on microhabitat temperature *T_m_* for individual species of *Erebia* butterflies. The fitted relationships are visualised in Fig. 1 and Fig. 2. Estimated d.f. describes the complexity of the fitted relationship; *e.d.f.* = 1 for a linear relationship, *e.d.f.* > 1 for a non-linear relationship.

Differences in body temperatures between the three low-altitude species/populations and the alpine species were significant based on a comparison of AIC of GAMs with and without the location as a grouping factor. The model of the dependence of *T_b_* on *T_a_* including the distinction between the low-altitude and the alpine sites had a lower AIC (Δ*AIC* = –382), differences between individual species included as a grouping factor were also highly significant (Δ*AIC* = –722). The same results hold for the dependence of *T_b_* on *T_m_* (Δ*AIC* = –266 for the effect of location and Δ*AIC* = –598 for the effect of species identity). Repeating the analysis with a subset of the data pertaining only to *E. euryale* yielded a significant difference between the low-altitude (*E. euryale*-CZ) and alpine (*E. euryale*-ALPS) populations (Δ*AIC* = –97 for the dependence of *T_b_* on *T_a_* and Δ*AIC* = –85 for the dependence of *T_b_* on *T_m_*).

Analysis of the thermoregulatory behaviour during sitting or nectaring showed that *E. aethiops* and *E. euryale*-CZ did not search for warm microclimates as much as the other species did (Fig. 3, Table 3). In *E. medusa,* the difference between microhabitat and air temperatures *T_m_* – *T_a_* during sitting and nectaring was more similar to the alpine species. Moreover, *E. medusa* explored microhabitat temperatures up to 32°C (Fig. 3C), the highest of all species studied. Excepting *E. aethiops, Erebia* species searched for warmer microclimates than the ambient air, i.e. *T_m_* > *T_a_* (Fig. 3). *E. aethiops* explored microhabitats which were similarly warm as the air (Fig. 3B), suggesting heat-avoidance behaviour. The high-altitude population of *E. euryale* searched for microhabitats on an average of 2.00 °C warmer than the air, while its low-altitude population was found in microhabitats on an average of only 0.64 °C warmer than the air.

**Figure 3.**
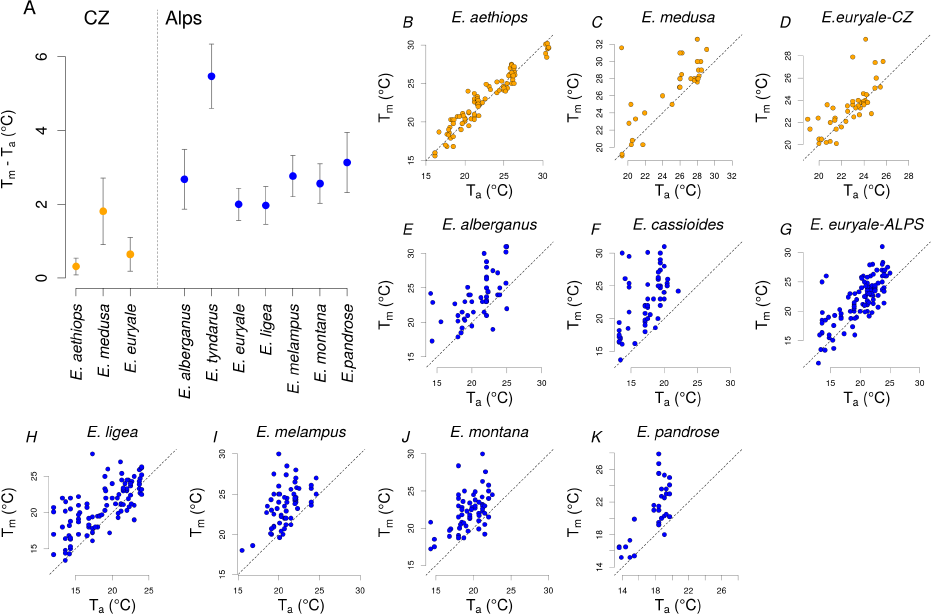
Behavioural thermoregulation is manifested by selectivity for microhabitats with temperature differing from the air temperature. Mean difference between microhabitat and air temperature *T_m_* — *T_a_* for individual species; vertical bars denote 95% confidence intervals (A). Data for settling and nectaring individuals of lowland (orange circles) and mountain (blue circles) *Erebia* butterflies are shown. The dependence of microhabitat temperature *T_m_* on air temperature *T_a_* for individual species and populations of lowland (B-D) and alpine (E-K) butterflies. Dotted lines show *T_m_* = *T_a_*.

The proportion of settling, nectaring or flying individuals was dependent on air temperature *T_a_* at least for one of these behavioural categories in *E. aethiops, E. euryale*-Alps, *E. ligea, E. montana* and *E. medusa* (Table 3, Fig. 4). Settling (i.e. various forms of basking behavior and resting) was the most frequent behaviour at low air temperatures (Fig. 4). The frequency of nectaring increased with increasing air temperatures in species from the Alps, but was hump-shaped in *E. aethiops* and *E. medusa.* The frequency of flying was nearly constant in all species, with the exception of *E. aethiops* which displayed an increase in the proportion of flying individuals with increasing air temperature.

**Figure 4.**
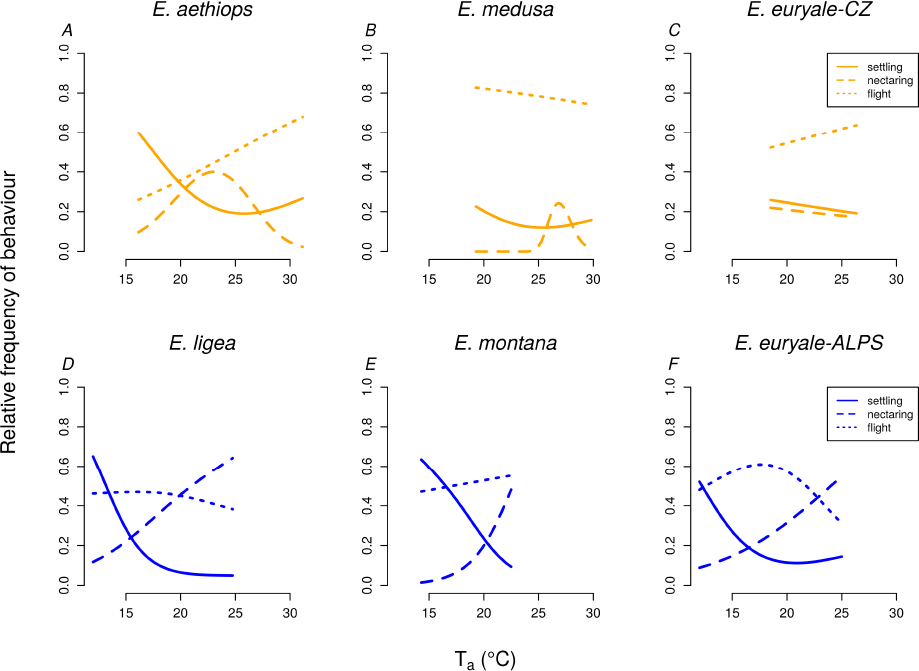
Main types of behaviour are affected by the air temperature. The dependence of the proportion of settling, nectaring and flying individuals on air temperature *T_a_* in individual species of *Erebia* butterflies as estimated by generalized additive models (GAM). Only species with at least some significant trends are displayed; full results of the GAMs are shown in Table 3.

**Table 3.**
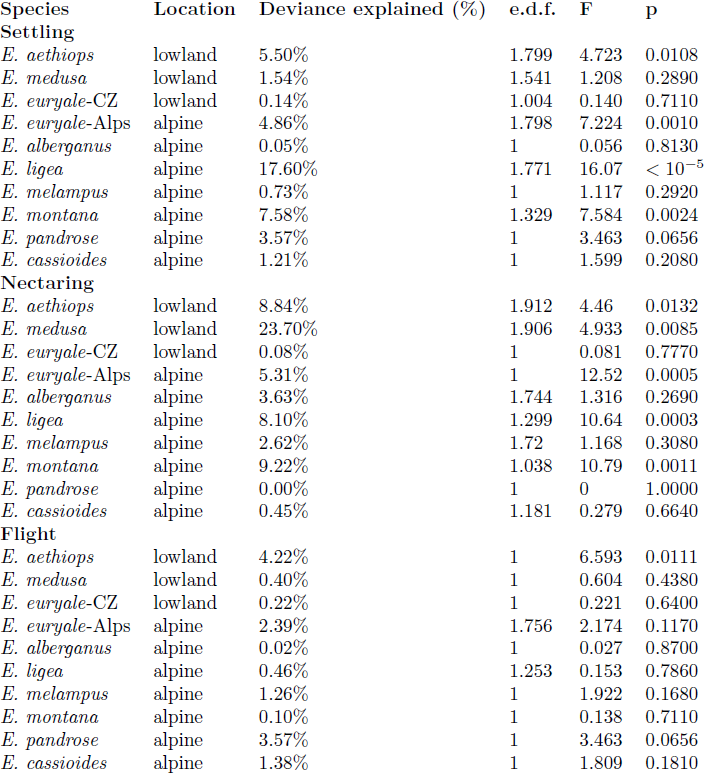
Proportion of settling, nectaring, and flying *Erebia* butterflies depends on air temperature. Results of generalized additive models (GAM) testing the dependence of the proportion of settling, nectaring and flying individuals of *Erebia* species on the air temperature *T_a_*. The fitted relationships are visualised in Fig. 4. Estimated d.f. describes the complexity of the fitted relationship; *e.d.f.* = 1 for a linear relationship, *e.d.f.* > 1 for a non-linear relationship.

| Species | Location | Deviance explained (%) | e.d.f. | F | p |
| --- | --- | --- | --- | --- | --- |
| **Settling** |  |  |  |  |  |
| *E. aethiops* | lowland | 5.50% | 1.799 | 4.723 | 0.0108 |
| *E. medusa* | lowland | 1.54% | 1.541 | 1.208 | 0.2890 |
| *E. euryale*-CZ | lowland | 0.14% | 1.004 | 0.140 | 0.7110 |
| *E. euryale*-Alps | alpine | 4.86% | 1.798 | 7.224 | 0.0010 |
| *E. alberganus* | alpine | 0.05% | 1 | 0.056 | 0.8130 |
| *E. ligea* | alpine | 17.60% | 1.771 | 16.07 | < 10<sup>-5</sup> |
| *E. melampus* | alpine | 0.73% | 1 | 1.117 | 0.2920 |
| *E. montana* | alpine | 7.58% | 1.329 | 7.584 | 0.0024 |
| *E. pandrose* | alpine | 3.57% | 1 | 3.463 | 0.0656 |
| *E. cassioides* | alpine | 1.21% | 1 | 1.599 | 0.2080 |
| **Nectaring** |  |  |  |  |  |
| *E. aethiops* | lowland | 8.84% | 1.912 | 4.46 | 0.0132 |
| *E. medusa* | lowland | 23.70% | 1.906 | 4.933 | 0.0085 |
| *E. euryale*-CZ | lowland | 0.08% | 1 | 0.081 | 0.7770 |
| *E. euryale*-Alps | alpine | 5.31% | 1 | 12.52 | 0.0005 |
| *E. alberganus* | alpine | 3.63% | 1.744 | 1.316 | 0.2690 |
| *E. ligea* | alpine | 8.10% | 1.299 | 10.64 | 0.0003 |
| *E. melampus* | alpine | 2.62% | 1.72 | 1.168 | 0.3080 |
| *E. montana* | alpine | 9.22% | 1.038 | 10.79 | 0.0011 |
| *E. pandrose* | alpine | 0.00% | 1 | 0 | 1.0000 |
| *E. cassioides* | alpine | 0.45% | 1.181 | 0.279 | 0.6640 |
| **Flight** |  |  |  |  |  |
| *E. aethiops* | lowland | 4.22% | 1 | 6.593 | 0.0111 |
| *E. medusa* | lowland | 0.40% | 1 | 0.604 | 0.4380 |
| *E. euryale*-CZ | lowland | 0.22% | 1 | 0.221 | 0.6400 |
| *E. euryale*-Alps | alpine | 2.39% | 1.756 | 2.174 | 0.1170 |
| *E. alberganus* | alpine | 0.02% | 1 | 0.027 | 0.8700 |
| *E. ligea* | alpine | 0.46% | 1.253 | 0.153 | 0.7860 |
| *E. melampus* | alpine | 1.26% | 1 | 1.922 | 0.1680 |
| *E. montana* | alpine | 0.10% | 1 | 0.138 | 0.7110 |
| *E. pandrose* | alpine | 3.57% | 1 | 3.463 | 0.0656 |
| *E. cassioides* | alpine | 1.38% | 1 | 1.809 | 0.1810 |

## Discussion

### Thermoregulatory strategies at thermal margins

Behaviour has the potential to modify physiological responses to evolutionary pressures such as the ongoing climate change [8, 44, 45]. Small ectothermic animals have to decrease their activity or rely on shade seeking behaviour to avoid overheating at the upper thermal margins of their range [10]. Interspecific differences in behavioural thermoregulation (e.g. species’ ability to explore colder microclimates in the woodland understorey) could modify the effect of extreme air temperatures and lead to different responses to external thermal constraints. Our results demonstrate that adults of low-altitude populations of *E. aethiops* and *E. medusa,* species of a mostly cold-dwelling butterfly genus, diversified in their thermoregulatory strategies and tolerance to high-temperatures (Fig. 2). We hypothesize that this differentiation of thermoregulatory strategies was driven by contrasting habitat preferences of these two species, similarly as has been observed for Mediterranean cicadas [46, 47], tropical lizards [21] or mountain species of the genus *Erebia* [14]. We also found differences between low-altitude and alpine populations of *E. euryale,* which probably reflect local adaptations to different climates. These local differences suggest that behavioural thermoregulation of adults is not able to buffer environmental pressures completely.

*E. aethiops,* which inhabits heterogeneous forest-steppe environments [18], did not actively enhance its body temperature and altered its activity in hot weather (see also [17]). *E. aethiops* was frequently observed flying in the shadow of trees during the warmest part of the day [17]. In contrast to other *Erebia* species studied, it showed a lack of selectivity for microhabitats warmer than the ambient air (Fig. 3). Furthermore, it was the only *Erebia* species in which the proportion of flying individuals increased at the highest air temperatures (Fig. 4). We observed them flying mostly among trees and shrubs (see also [17]), which may serve to decrease body temperature, although there may be a range of other factors shaping the activity pattern, such as behaviour related to reproduction. *E. aethiops* mainly settled on vegetation during the lowest air temperatures experienced (15-20<u>º</u>C). However, these air temperatures are suitable for activity for the other *Erebia* species; alpine relatives were found in flight at temperatures as low as 12<u>º</u>C [14]. Settling during temperatures potentially suitable for activity may be conditioned by resting in a colder microclimate under shrubs and trees [17] and by a lower effort to warm up in this species. *E. aethiops* displayed a shift of activity to higher temperatures than other species also in nectaring (Fig. 4).

Contrary to *E. aethiops, E. medusa,* a species of more open grasslands, effectively heated up under low air temperature and linearly enhanced its body temperature up to 39°C; i.e., higher than other *Erebia* species (see also [14]). These differences illustrate that species with an ancestral cold-climate preference [32] have the potential to adjust to environmental conditions in warm lowlands and that behavioural traits, such as habitat use, determine species’ thermal preferences and may affect their responses to predicted climate warming. Indeed, species living in cold climates generally seem to retain relatively high upper thermal tolerance limits, and they differ from species living in warmer climates mostly by their unique tolerance to cold [48]. In agreement, our data further show that the body temperatures of *E. aethiops* and the low-altitude population of *E. euryale* differed from the other species mostly under low microhabitat temperatures, while they were similar under higher temperatures (Fig. 1E-H, Fig. 2B), which points to different physiological responses to cold rather than to heat.

Both lowland species experienced higher maximal air temperatures (around 30<u>º</u>C) than their mountain congeners (Fig. 4) because of differences in local climates. It seems that the extreme temperatures limited nectaring of the two lowland species. *E. aethiops* reached the peak of its nectaring activity around 24<u>º</u>C and *E. medusa* around 26<u>º</u>C. The decrease of nectaring activities of the lowland species during the warmest periods of the day, not observed in the mountain species, could be caused by desiccation of nectar or by the need of the butterflies to prevent overheating. Thus, the timing of diurnal activities may contribute to the *Erebia* thermoregulation together with microhabitat choice [17, 49] and facilitate species survival on their range margins. However, laboratory measurements of thermal tolerance traits will be necessary to properly evaluate the effect of thermal extremes on different species.

### Behavioural thermoregulation and evolutionary adaptation to climate change

Long-term persistence in a changing climate requires adaptations such as adjustments of physiological mechanisms responsible for thermal tolerance. It is possible that behavioural thermoregulation, which is a potent short-term solution for dealing with a changing climate, constrains the adaptive potential of behaviourally plastic species by buffering the selective pressure of temperature [22]. This would lead to a limited potential to adapt to climate change by evolution of physiological traits [8].This phenomenon has been termed the “Bogert effect” and supporting evidence comes from lizards [8, 22, 23] as well as invertebrates, e.g. *Drosophila* [50], although other studies dispute its general applicability [51]. However, there is a large amount of evidence that behavioural plasticity, and phenotypic plasticity in general, can drive rather than prevent evolutionary changes (reviewed in [52,53]). The effects of behavioural plasticity on evolution are likely context-dependent and may differ between different traits. Adults of *Erebia* butterflies thermoregulate behaviourally but their thermoregulatory strategies and preferred body temperatures are not conserved according to our data; i.e. individual species have different requirements, which does not provide support for the Bogert effect in *Erebia*.

According to our measurements, responses to temperature also differed between populations of *E. euryale* from two sites differing in climate. The low-altitude population *E. euryale*-CZ (900m a.s.l.) kept lower body temperatures than the high-altitude population *E. euryale*-Alps (1800m a.s.l.) at comparable ambient temperatures (Fig. 2). This suggests that the lowland population is more "lazy" in warming up and its activity is constrained to periods of the day with optimal air temperature. In contrast, mountain species are limited by a shorter time window of favourable temperatures, so they need to be active whenever possible. In a similar case in grasshoppers, Samietz et al. [54] found that individuals increased their body temperatures via mobility and basking more at higher altitudes and demonstrated that this was due to local adaptation. Evidence of local adaptation in thermoregulation was found also in butterflies [55, 56], lizards [21, 44, 57], and other taxa. An alternative explanation for the observed differences between *E. euryale* populations is phenotypic plasticity. Resolving the relative contributions of these interacting factors requires transplant or laboratory experiments [54, 58]. It is also important to note that main selective agents driving the evolution of thermoregulatory strategies are temperature extremes, rather than long-term means, and that maximal temperatures seem to be more limiting than minimal temperatures in terrestrial ectotherms [59]. Also, there is mounting evidence that tolerance to heat does not evolve as easily as tolerance to cold [44, 48]. Moreover, the effect of temperature may differ across developmental stages, which needs to be investigated [60].

### Habitat heterogeneity and species survival in a changing climate

Habitat heterogeneity provides opportunities for species of ectotherms with flexible behaviour, such as *Erebia* butterflies studied here, to find a suitable microclimate and to obtain the resources they require. The butterflies can adjust their microhabitat use and shift the timing of various activities. Thus, habitat heterogeneity may be a key to their survival in a changing climate [61]. Mountain species may explore warmer microhabitats to enhance their body temperature [14], while lowland species have the possibility to retreat to the shade in case the weather gets too hot. Whereas mountain landscapes provide heterogeneity by their rugged terrain, lowlands are more prone to landscape homogenization and suffer from higher human pressure [62]. Lowland species living in woodlands, such as *E. aethiops* [18], may thus be more endangered by a combination of landscape homogenization and climate change than alpine species or species of open grasslands, such as *E. medusa.* They might be both more sensitive to high temperatures because of their lower preferred temperatures [14] and have limited opportunities for microhabitat shifts, e.g. more intensive shade seeking would be difficult because they already inhabit relatively cold microhabitats [21]. On the other hand, it could be expected that under climate warming, a grassland species such as *E. medusa* would retreat to woodlands which provide shade. Indeed, this species inhabits such habitats in some parts of its range [39]. It remains to be tested whether its local habitat preferences correlate with the local climate.

It is important to highlight that there are other factors associated with climate change, such as changes in precipitation and snow cover. These factors may modify the effects of temperature and affect different developmental stages. Our study focused only on adults, without considering eggs, larvae, and pupae. It is already known, however, that the effect of temperature may differ across developmental stages [60], which needs to be investigated further. Environmental conditions experienced by larvae are to a certain degree predetermined by adult behaviour. For example, in *Euphydryas editha,* differences in adult phenology and egg placement by ovipositioning females lead to differences in the thermal environment experienced by larvae [13]. Not only adults, but also larvae can thermoregulate behaviourally [63], but not during the winter when they are dormant. Snow provides insulation during overwintering, and thus affects the survival of dormant stages. Decreasing snow cover may thus adversely affect butterflies by increasing frost-related larval mortality during overwintering [29, 35, 64]. Data on tolerances of all stages to temperature and other key environmental factors are thus necessary to reveal the main selective pressures [45, 65, 66] and to better understand the often varied responses of species to climate change [67].

## Supporting Information

### S1 File

#### Raw data

The file contains measurements of body temperature, air temperature, microhabitat temperature, and behaviour prior to capture of *Erebia aethiops, E. euryale*, and *E. medusa* studied in the Czech Republic.

## Acknowledgements

We would like to thank D. Novotný and J. Peltanová for help with data collection in the field and to M. Konvička for inspiring discussions on butterfly ecology and conservation. Both JK and IK are supported by the Czech Science Foundation (projects GP14-10035P and GA14-33733S). We also thank two reviewers for their thoughtful comments and Matthew Sweney for proofreading the manuscript.

## References

1. Konvicka M, Maradova M, Benes J, Fric Z, Kepka P. Uphill shifts in distribution of butterflies in the Czech Republic: effects of changing climate detected on a regional scale. Global Ecology and Biogeography. 2003;12(5):403–410.

2. Roth T, Plattner M, Amrhein V. Plants, birds and butterflies: short-term responses of species communities to climate warming vary by taxon and with altitude. PLOS One. 2014;9(1):e82490.

3. Karl I, Schmitt T, Fischer K. Genetic differentiation between alpine and lowland populations of a butterfly is related to PGI enzyme genotype. Ecography. 2009;32(3):488–496.

4. Kjaersgaard A, Blanckenhorn WU, Pertoldi C, Loeschcke V, Kaufmann C, Hald B, et al. Plasticity in behavioural responses and resistance to temperature stress in *Musca domestica*. Animal Behaviour. 2015;99:123–130.

5. Chown SL, Hoffmann AA, Kristensen TN, Angilletta Jr MJ, Stenseth NC, Pertoldi C. Adapting to climate change: a perspective from evolutionary physiology. Climate Research. 2010;43(1):3–15.

6. Vila R, Bell CD, Macniven R, Goldman-Huertas B, Ree RH, Marshall CR, et al. Phylogeny and palaeoecology of *Polyommatus* blue butterflies show Beringia was a climate-regulated gateway to the New World. Proceedings of the Royal Society of London B: Biological Sciences. 2011;p. 20102213.

7. Kleckova I, Cesanek M, Fric Z, Pellissier L. Diversification of the cold-adapted butterfly genus *Oeneis* related to Holarctic biogeography and climatic niche shifts. Molecular Phylogenetics and Evolution. 2015;92:255–265.

8. Buckley LB, Ehrenberger JC, Angilletta MJ. Thermoregulatory behaviour limits local adaptation of thermal niches and confers sensitivity to climate change. Functional Ecology. 2015;29(8):1038–1047.

9. Illân JG, Gutierrez D, Diez SB, Wilson RJ. Elevational trends in butterfly phenology: implications for species responses to climate change. Ecological Entomology. 2012;37(2):134–144.

10. Sunday JM, Bates AE, Kearney MR, Colwell RK, Dulvy NK, Longino JT, et al. Thermal-safety margins and the necessity of thermoregulatory behavior across latitude and elevation. Proceedings of the National Academy of Sciences. 2014;111(15):5610–5615.

11. Turlure C, Choutt J, Baguette M, Van Dyck H. Microclimatic buffering and resource-based habitat in a glacial relict butterfly: significance for conservation under climate change. Global Change Biology. 2010;16(6):1883–1893.

12. Lawson CR, Bennie JJ, Thomas CD, Hodgson JA, Wilson RJ. Local and landscape management of an expanding range margin under climate change. Journal of Applied Ecology. 2012;49(3):552–561.

13. Bennett NL, Severns PM, Parmesan C, Singer MC. Geographic mosaics of phenology, host preference, adult size and microhabitat choice predict butterfly resilience to climate warming. Oikos. 2015;124(1):41–53.

14. Kleckova I, Konvicka M, Klecka J. Thermoregulation and microhabitat use in mountain butterflies of the genus *Erebia:* importance of fine-scale habitat heterogeneity. Journal of Thermal Biology. 2014;41:50–58.

15. Kearney M, Shine R, Porter WP. The potential for behavioral thermoregulation to buffer “cold-blooded” animals against climate warming. Proceedings of the National Academy of Sciences. 2009;106(10):3835–3840.

16. Marschalek DA, Klein Sr MW. Distribution, ecology, and conservation of Hermes copper (Lycaenidae: *Lycaena [Hermelycaena] hermes*). Journal of Insect Conservation. 2010;14(6):721–730.

17. Slamova I, Klecka J, Konvicka M. Diurnal behavior and habitat preferences of *Erebia aethiops,* an aberrant lowland species of a mountain butterfly clade. Journal of Insect Behavior. 2011; 24(3):230–246.

18. Slamova I, Klecka J, Konvicka M. Woodland and grassland mosaic from a butterfly perspective: habitat use by *Erebia aethiops* (Lepidoptera: Satyridae). Insect Conservation and Diversity. 2013;6(3):243–254.

19. Parmesan C, Ryrholm N, Stefanescu C, Hill JK, Thomas CD, Descimon H, et al. Poleward shifts in geographical ranges of butterfly species associated with regional warming. Nature. 1999;399(6736):579–583.

20. Habel J, Ivinskis P, Schmitt T. On the limit of altitudinal range shifts-Population genetics of relict butterfly populations. Acta Zoologica Academiae Scientiarum Hungaricae. 2010;56(4):383–393.

21. Huey RB, Deutsch CA, Tewksbury JJ, Vitt LJ, Hertz PE, Perez HJA, et al. Why tropical forest lizards are vulnerable to climate warming. Proceedings of the Royal Society of London B: Biological Sciences. 2009;p. 20081957.

22. Bogert CM. Thermoregulation in reptiles, a factor in evolution. Evolution. 1949;3(3):195–211.

23. Huey RB, Hertz PE, Sinervo B. Behavioral drive versus behavioral inertia in evolution: a null model approach. American Naturalist. 2003;161(3):357–366.

24. Wilson RJ, Gutierrez D, Gutierrez J, Martinez D, Agudo R, Monserrat VJ. Changes to the elevational limits and extent of species ranges associated with climate change. Ecology Letters. 2005;8(11):1138–1146.

25. Franzen M, Molander M. How threatened are alpine environments? A cross taxonomic study. Biodiversity and Conservation. 2012;21(2):517–526.

26. Karl I, Stoks R, De Block M, Janowitz SA, Fischer K. Temperature extremes and butterfly fitness: conflicting evidence from life history and immune function. Global Change Biology. 2011;17(2):676–687.

27. Pellissier L, Brathen KA, Vittoz P, Yoccoz NG, Dubuis A, Meier ES, et al. Thermal niches are more conserved at cold than warm limits in arctic-alpine plant species. Global Ecology and Biogeography. 2013;22(8):933–941.

28. Ashton S, Gutierrez D, Wilson RJ. Effects of temperature and elevation on habitat use by a rare mountain butterfly: implications for species responses to climate change. Ecological Entomology. 2009;34(4):437–446.

29. Vrba P, Konvicka M, Nedved O. Reverse altitudinal cline in cold hardiness among *Erebia* butterflies. CryoLetters. 2012;33(4):251–258.

30. Louy D, Habel JC, Ulrich W, Schmitt T. Out of the Alps: The Biogeography of a disjunctly distributed mountain butterfly, the Almond-eyed ringlet *Erebia alberganus* (Lepidoptera, Satyrinae). Journal of Heredity. 2013;p. est081.

31. Besold J, Schmitt T. More northern than ever thought: refugia of the Woodland Ringlet butterfly *Erebia medusa* (Nymphalidae: Satyrinae) in Northern Central Europe. Journal of Zoological Systematics and Evolutionary Research. 2015;53(1):67–75.

32. Pena C, Witthauer H, Kleckova I, Fric Z, Wahlberg N. Adaptive radiations in butterflies: evolutionary history of the genus *Erebia* (Nymphalidae: Satyrinae). Biological Journal of the Linnean Society. 2015;116:449–467.

33. Schmitt T, Hewitt G. The genetic pattern of population threat and loss: a case study of butterflies. Molecular Ecology. 2004;13(1):21–31.

34. De Groot M, Rebusek F, Grobelnik V, Govedic M, Salamun A, Verovnik R. Distribution modelling as an approach to the conservation of a threatened alpine endemic butterfly (Lepidoptera: Satyridae). European Journal of Entomology. 2009;106(1):77–84.

35. Scalercio S, Bonacci T, Mazzei A, Pizzolotto R, Brandmayr P. Better up, worse down: bidirectional consequences of three decades of climate change on a relict population of *Erebia cassioides*. Journal of Insect Conservation. 2014;18(4):643–650.

36. Stuhldreher G, Fartmann T. When habitat management can be a bad thing: effects of habitat quality, isolation and climate on a declining grassland butterfly. Journal of Insect Conservation. 2014;18(5):965–979.

37. Sonderegger P. Die Erebien der Schweiz: (Lepidoptera: Satyrinae, Genus *Erebia*). Eigenverlag; 2005.

38. van Swaay C, Cuttelod A, Collins S, Maes D, López Munguira M, Šašić M, et al. European red list of butterflies. Publications Office of the European Union, Luxembourg; 2010.

39. Beneš J, Konvička M, Dvořák J, Fric Z, Havelda Z, Pavlíčko A, et al. Motýli České republiky: Rozšíření a ochrana I., II. (Butterflies of the Czech Republic: Distribution and conservation I., II.). SOM, Prague; 2002.

40. van Swaay C, Warren M. Red data book of European butterflies (Rhopalocera). vol. 99. Council of Europe; 1999.

41. Stuhldreher G, Hermann G, Fartmann T. Cold-adapted species in a warming world-an explorative study on the impact of high winter temperatures on a continental butterfly. Entomologia Experimentalis et Applicata. 2014;151(3):270–279.

42. R Core Team. R: A Language and Environment for Statistical Computing. Vienna, Austria; 2013. Available from: http://www.R-project.org/.

43. Wood S. Generalized additive models: an introduction with R. CRC press; 2006.

44. Munoz MM, Stimola MA, Algar AC, Conover A, Rodriguez AJ, Landestoy MA, et al. Evolutionary stasis and lability in thermal physiology in a group of tropical lizards. Proceedings of the Royal Society of London B: Biological Sciences. 2014;281(1778):20132433.

45. Gunderson AR, Stillman JH. Plasticity in thermal tolerance has limited potential to buffer ectotherms from global warming. Proceedings of the Royal Society of London B: Biological Sciences. 2015;282(1808):20150401.

46. Sanborn AF, Heath JE, Phillips PK, Heath MS, Noriega FG. Thermal Adaptation and Diversity in Tropical Ecosystems: Evidence from Cicadas (Hemiptera, Cicadidae). PLOS One. 2011;6(12):e29368.

47. Sanborn AF, Phillips PK, Heath JE, Heath MS. Comparative thermal adaptation in cicadas (Hemiptera: Cicadidae) inhabiting Mediterranean ecosystems. Journal of Thermal Biology. 2011;36(2):150–155.

48. Araùjo MB, Ferri-Yánez F, Bozinovic F, Marquet PA, Valladares F, Chown SL. Heat freezes niche evolution. Ecology Letters. 2013;16(9):1206–1219.

49. Konvicka M, Benes J, Kuras T. Microdistribution and diurnal behaviour of two sympatric mountain butterflies *(Erebia epiphron* and *E. euryale*): relations to vegetation and weather. Biologia. 2002;57(2):223–233.

50. Castaneda LE, Balanya J, Rezende EL, Santos M. Vanishing chromosomal inversion clines in *Drosophila subobscura* from Chile: is behavioral thermoregulation to blame? American Naturalist. 2013;182(2):249–259.

51. Mitchell KA, Sinclair BJ, Terblanche JS. Ontogenetic variation in cold tolerance plasticity in *Drosophila:* is the Bogert effect bogus? Naturwissenschaften. 2013;100(3):281–284.

52. Foster SA. Evolution of behavioural phenotypes: influences of ancestry and expression. Animal Behaviour. 2013;85(5):1061–1075.

53. Zuk M, Bastiaans E, Langkilde T, Swanger E. The role of behaviour in the establishment of novel traits. Animal Behaviour. 2014;92:333–344.

54. Samietz J, Salser M, Dingle H. Altitudinal variation in behavioural thermoregulation: local adaptation vs. plasticity in California grasshoppers. Journal of Evolutionary Biology. 2005;18(4):1087–1096.

55. Ellers J, Boggs CL. Functional ecological implications of intraspecific differences in wing melanization in *Colias* butterflies. Biological Journal of the Linnean Society. 2004;82(1):79–87.

56. Pitteloud C, Arrigo N, Suchan T, Mastretta-Yanes A, Vila R, Dinca V, et al. Ecological character displacement and geographical context of lineage divergence: macro-evolutionary insights in the butterfly genus *Pyrgus*. Submitted;.

57. Moritz C, Langham G, Kearney M, Krockenberger A, VanDerWal J, Williams S. Integrating phylogeography and physiology reveals divergence of thermal traits between central and peripheral lineages of tropical rainforest lizards. Philosophical Transactions of the Royal Society B: Biological Sciences. 2012;367(1596):1680–1687.

58. Merckx T, Van Dyck H. Landscape structure and phenotypic plasticity in flight morphology in the butterfly *Pararge aegeria*. Oikos. 2006;113(2):226–232.

59. Hoffmann AA, Chown SL, Clusella-Trullas S. Upper thermal limits in terrestrial ectotherms: how constrained are they? Functional Ecology. 2013;27(4):934–949.

60. Turlure C, Van Dyck H, Goffart P, Schtickzelle N, et al. Resource-based habitat use in *Lycaena helle*: Significance of a functional, ecological niche-oriented approach. In: Habel J, Meyer M, Schmitt T, editors. Jewels in the mist: A synopsis on the endangered Violet copper butterfly, *Lycaena helle*. Pensoft Publishers; 2014. p. 67–86.

61. Lawson CR, Bennie JJ, Thomas CD, Hodgson JA, Wilson RJ. Active management of protected areas enhances metapopulation expansion under climate change. Conservation Letters. 2014;7(2):111–118.

62. Vellend M, Verheyen K, Flinn KM, Jacquemyn H, Kolb A, Van Calster H, et al. Homogenization of forest plant communities and weakening of species-environment relationships via agricultural land use. Journal of Ecology. 2007;95(3):565–573.

63. Sherman PW, Watt WB. The thermal ecology of some *Colias* butterfly larvae. Journal of Comparative Physiology. 1973;83(1):25–40.

64. Lamb R, Turnock W, Hayhoe H. Winter survival and outbreaks of bertha armyworm, *Mamestra configurata* (Lepidoptera: Noctuidae), on canola. The Canadian Entomologist. 1985;117(06):727–736.

65. Mercader R, Scriber J. Asymmetrical thermal constraints on the parapatric species boundaries of two widespread generalist butterflies. Ecological Entomology. 2008;33(4):537–545.

66. Radchuk V, Turlure C, Schtickzelle N. Each life stage matters: the importance of assessing the response to climate change over the complete life cycle in butterflies. Journal of Animal Ecology. 2013;82(1):275–285.

67. Parmesan C. Ecological and evolutionary responses to recent climate change. Annual Review of Ecology, Evolution, and Systematics. 2006;37:637–669.

